# Opposing effects of histone H2A.Z on memory, transcription and pathology in male and female Alzheimer’s disease mice and patients

**DOI:** 10.1101/2025.05.28.656659

**Authors:** Jian Qi Luo, Luca A. Hategan, Samantha Creighton, Shan Hua, Timothy A.B. McLean, Tarkan Ahmad Dahi, Zhenhong Jin, Fardad Pirri, Stephen M Winston, Isaiah L Reeves, Margaret Fahnestock, Mark A. Brimble, Brandon J. Walters, Patrick J. Murphy, Iva B. Zovkic

## Abstract

Alzheimer’s disease (AD) is a devastating neurodegenerative disease that disproportionately impacts women, but underlying mechanisms for sex-divergent outcomes are unknown. Here, we show that the histone variant H2A.Z is a novel sex-specific regulator of AD in human patients and AD model mice. Specifically, H2A.Z binding in chromatin declines in female and increases in male AD patients, indicating opposite patterns of AD-related H2A.Z dysregulation each sex. These sex differences were recapitulated in the 5xFAD model of AD, in which females accumulated H2A.Z at early disease stages and lost H2A.Z as the disease progressed, suggesting that H2A.Z occupancy shifts with advancing disease. Males showed no change in H2A.Z binding in early disease, but exhibited increased binding as disease progressed, albeit to a lesser extent than females. Consistent with sex-specific H2A.Z dysregulation, H2A.Z depletion produced sex-specific changes in gene expression, whereby H2A.Z was more repressive in female than in male mice and in 5xFAD than in WT males, suggesting that H2A.Z’s role in transcription varies with sex and disease. Moreover, H2A.Z depletion improved memory and AD pathology in females, while impairing memory and worsening pathology in male mice. Together, these data suggest that H2A.Z is protective in males and detrimental in females with AD, with key implications for sex-specific therapeutic targeting of chromatin factors.

## Introduction

Alzheimer’s disease (AD) is a devastating neurodegenerative disease that disproportionately impacts women, who account for two thirds of AD patients [1]. Growing evidence suggests that therapeutic outcomes are sex-biased, with drugs like lecanemab exhibiting better efficacy in male than female patients [2]. Such outcomes are consistent with growing evidence for sex differences in underlying disease pathways and evidence for sex-specific risk genes for AD [3, 4], but mechanisms underlying sex differences in AD are understudied.

Epigenetic factors are promising candidates for disease regulation because they integrate information from upstream signals and translate them into downstream changes in gene expression that impact all aspects of cell function and disease progression. Moreover, epigenetic factors may promote sex-specific disease outcomes because of extensive sex differences in disease- and age-related changes in gene expression [3, 5, 6]. Although DNA methylation and histone post-translational modifications (PTMs) have established roles in AD, a role for histone variants has not been studied despite these histones comprising the majority of adult neuronal chromatin. Specifically, histone variants are functionally and structurally distinct histone types that replace canonical histones in chromatin in response to neuronal activity and are critical for normal neural plasticity and memory formation [7–21]. Canonical histones are coupled to DNA replication and are produced primarily during cell division, whereas histone variants are independent of replication and thus become the primary source of histones in post-mitotic neurons [14, 22]. This over-representation of histone variants in the aging brain is indicative of their disproportionate contribution to age-related diseases, but their role in AD has never been studied.

The histone variant H2A.Z is of particular interest because we previously reported that H2A.Z depletion has sex-specific effects on memory, its transcription and its binding are regulated by sex hormones, and it is required for androgen-mediated memory regulation [9–14]. In addition, we found a suppressive role of H2A.Z in memory [7–12] and showed that its levels accumulate with aging in male mice [14], indicating that altered regulation of this histone may contribute to AD-related memory decline.

Here, we report that H2A.Z is regulated in a sex-specific manner in human AD patients and AD model mice, whereby H2A.Z binding declines in female and increases in male AD patients. Moreover, shifts in H2A.Z binding are modified with disease progression and result in sex-specific disease outcomes, whereby H2A.Z depletion improves memory and AD pathology in females, while impairing memory and worsening pathology in males. These data establish histone variants as novel regulators of AD and suggest that sex-specific dysregulation of AD contributes to sex differences in disease risk and progression.

## Methods

### Animals

Male and female 5xFAD mice were purchased from Jackson Labs and bred in our colony with C57BL/6J mice to create heterozygous 5xFAD mice. Mice were housed in groups on a 12h light cycle (lights on at 8 am) with *ad libitum* access to food and water. All procedures were approved by the University of Toronto Animal Care Committee and complied with institutional guidelines and the Canadian Council on Animal Care.

### Chromatin immunoprecipitation (ChIP)

Post-mortem hippocampal tissue was donated by the Douglas-Bell Canada Brain Bank. Human post-mortem brain samples were first sectioned into ∼25 mg pieces on dry ice and stored at –80°C until processing. For cross-linking, samples were incubated with 1% formaldehyde containing a cocktail of protease inhibitors (Cell Signaling Technology, #5871) for 5 minutes at room temperature (RT), followed by quenching with 125 mM glycine for 5 minutes at RT. Tissues were then washed six times with 200 µL of ice-cold PBS. Samples were homogenized on ice using pellet pestles in 300 µL of SDS lysis buffer (50 mM Tris-HCl, pH 8.1; 10 mM EDTA; 1% SDS; and protease inhibitors). Lysates were incubated on ice for 10 minutes to ensure complete lysis. Chromatin was initially sheared using a probe sonicator for 10 seconds at 40% power, followed by further fragmentation using the Bioruptor (Diagenode) with 50 cycles of 30 seconds ON/30 seconds OFF at full power. Samples were vortexed and briefly centrifuged every 10 cycles to ensure consistent shearing. Lysates were centrifuged at 17,000×g for 5 minutes at 4°C to remove insoluble debris. The supernatant was split into two aliquots for input and immunoprecipitation (IP). A 45 µL aliquot (15% of the lysate) was saved as input, and the remaining lysate was diluted 1:8 with ChIP dilution buffer (0.01% SDS, 1.1% Triton X-100, 1.2 mM EDTA, 16.7 mM Tris-HCl, pH 8.1, 167 mM NaCl) for IP. Magna ChIP Protein G magnetic beads (Millipore, #16-662) were blocked in 0.5% BSA-PBS for 5 minutes on a rotator at RT, repeated three times. Blocked beads were washed twice with ChIP dilution buffer and resuspended to the original volume. Diluted chromatin was incubated with 50 µL of blocked beads and 3 µL of H2A.Z antibody (Millipore, ABE1348), and rotated overnight at 4°C. The next day, beads were collected using a magnetic separator and washed sequentially with 1 mL of the following buffers: Low-salt buffer (0.1% SDS, 1% Triton X-100, 2 mM EDTA, 20 mM Tris-HCl, pH 8.1, 150 mM NaCl), High-salt buffer (0.1% SDS, 1% Triton X-100, 2 mM EDTA, 20 mM Tris-HCl, pH 8.1, 500 mM NaCl), LiCl buffer (0.25 M LiCl, 1% IGEPAL CA-630, 1% sodium deoxycholate, 1 mM EDTA, 10 mM Tris-HCl, pH 8.1), TE buffer (10 mM Tris-HCl, pH 8.0, 1 mM EDTA). For each wash, beads were resuspended by gentle inversion and rotated at 4°C for 5 minutes. Immune complexes were eluted by incubating beads in 90 µL of 1× TE buffer containing 6.6 µg of RNase A (Thermo Fisher, EN0531) at 37°C for 30 minutes. Subsequently, 10 µL of 10% SDS (final concentration 1%) and 1 µL of Proteinase K (Roche, 3115828001) were added, followed by incubation at 65°C for 2 hours and then 95°C for 10 minutes. Input samples were treated with 3× the volume of TE-RNase buffer/SDS/1.5ul of Pk and processed similarly. DNA was purified using the Monarch® PCR & DNA Cleanup Kit (NEB, 5 µg), following the manufacturer’s protocol with the modification of using pre-warmed (50°C) elution buffer. To prevent column clogging, input samples were split into two tubes for purification and subsequently pooled before library preparation.

To ensure sufficient starting material for library construction, two rounds of immunoprecipitation were performed, each using 25 mg of tissue. DNA from both rounds was pooled and submitted for library preparation at the Princess Margaret Genomics Centre. Libraries were sequenced on two 1.5B flow cells using paired-end 50 bp (PE50) reads, yielding approximately 80 million reads per library.

The ChIP-seq protocol for mouse brain tissue followed the human ChIP protocol with several modifications. Lysates were sonicated for 40 cycles using the Bioruptor. Decrosslinked DNA was purified using the QIAquick PCR Purification Kit (Qiagen, Cat# 28104). Only a single round of IP was performed before library preparation.

DNA concentrations of ChIP and input samples were measured using a Qubit 3.0 fluorometer. For library construction, 5–10 ng of ChIP DNA or 100 ng of input DNA was used with the NEBNext® Ultra™ II DNA Library Prep Kit for Illumina® (E7645S), following the manufacturer’s instructions with the exception of omitting size selection for input DNA. Samples were barcoded using NEBNext® Multiplex Oligos for Illumina® (96 Unique Dual Index Primer Pairs, E6440), and bead cleanup was performed using the NucleoMag NGS Clean-up and Size Select Kit (Macherey-Nagel, MN-744970.50). Final libraries were quantified with Qubit and assessed for size distribution using a Bioanalyzer. Libraries were sequenced on a 1.5B flow cell using PE50 reads, generating ∼50 million reads per library.

### Bioinformatics

#### ChIP-seq

Reads in Fastq format were quality controlled using fastqc (version 0.11.9). Mouse samples were aligned to the *Mus musculus* mm10 genome (Ensembl) and human samples were aligned to the *Homo sapiens* hg38 genome (Ensembl) using bowtie-2 (version 2.5.1) with paired-end settings. Adapters were trimmed prior to alignment using trimmomatic (version 0.39). Raw SAM files were converted to BAM using samtools (version 1.12) and deduplicated to retain only uniquely mapped paired reads at Q greater than 30. Filtered BAM files were imported into R (version 4.3.1) and processed using the csaw (version 1.36.1). Briefly, quality control was conducted using ChIPQC (version 1.37.0) and cross-correlation analysis was conducted in csaw. Counting was performed using the windowCounts function in csaw with a width parameter set to 150bp to cover the length of the nucleosome [23–25]. A background width of 10kb was set in the windowCounts function in csaw to compute the background signal with the bin flag set to true. Counts were filtered using the filterWindowsGlobal function in csaw, and reads passing a log_2_ [23] enrichment over background were retained in downstream analyses. Filtered counts were analyzed with MA plots, and count matrices were normalized either on the background binned read counts, or offsets were applied where trended biases were observed. Differential binding analysis was conducted in edgeR (version 3.42.4) by importing the retained counts matrix and normalization coefficients generated with csaw. When applicable, samples were batch corrected by modelling the batch coefficients in edgeR. All counts were merged to 500bp to a tolerance of 100bp in csaw using the mergeResults function in csaw. Volcano plots were generated in ggplot2 (version 3.4.2) from the merged counts, relating the representative log fold change and the -log10(FDR). A log fold change cutoff of 1, and an FDR cutoff of 0.05 were set [23]. Merged counts were annotated for closest gene and position along the gene body using the detailRanges function in R, passing the species specific TxDb or OrgDb objects, respectively (TxDb.Mmusculus.UCSC.mm10.knownGene version 3.10.0, org.Mm.eg.db version 3.17.0, TxDb.Hsapiens.UCSC.hg38.knownGene version 3.17.0, org.Hs.eg.db version 3.17.0). All other settings were left as defaults within csaw.

#### TSS heat plots

To generate representative profile plots around the TSS, within group, filtered, deduplicated BAM files were merged using samtools. Merged BAM files were converted to the BigWig format using deepTools (version 3.5.1). Using the computeMatrix function, with the reference-point argument, scores per genome region were calculated +/- 2000bp around the TSS. GTF files from either species were provided, respectively (Ensembl). To plot the heat map, plotHeatmap was called on the output matrix in deepTools using default settings.

#### Pie charts

To ascertain the location of H2A.Z binding throughout either the human or mouse genome, respectively, all merged counts from each comparison group, respectively, were processed to the BED format using a custom R script. Additionally, BED files were subset based on their significance scores, creating sites that were depleted of H2A.Z (FDR < 0.05 and log fold change<-1) and accumulated H2A.Z (FDR<0.05 and logFC > 1) per condition, respectively. BED files were annotated using annotatePeaks in Homer (version 5.1) using default settings.

#### H2A.Z enrichment on histone PTMs

To visualize H2A.Z binding at key histone PTMs, H2A.Z binding was assessed at differential histone PTMs site using a set of custom R scripts. Briefly, differential peaks of key histone PTMs from Gjoneska et al. [26] were separated into their identities (H3K4me3, H3K27Ac, and H3K27Me3) and significance (accumulating *vs* depleting) from the differential peak file [26]. Peak sites were lifted over to the mm10 annotations using the mm9ToMm10 chain file (Ensembl) and the liftOver function in R (rtracklayer, version 1.61.0). BED files were generated with these peaks using a custom R script, and profile plots of H2A.Z at these loci were generated using deepTools. Briefly, within group merged BigWig were used. computeMatrix with the reference-point argument set to center, in deepTools was called on the key histone PTM peaks as BED files, computing scores per genome region at these peak centers (+/- 2000bp). plotProfile was called to visualize the H2A.Z signal at these sites, partitioning signal based on genotype.

#### Log-log plots

To compare the change in H2A.Z binding over time, plots relating the log fold change of the two time points were generated using a custom R script. Briefly, merged counts were first preprocessed to track changes at the same loci over time. The overlapping coordinates at each time point were determined using the overlapsAny function in R (GenomicRanges version 1.52.0), these overlapping regions were then subset and the log fold change at each time point were plotted using ggplot2. A linear regression was fit to these plots in R.

### RNA sequencing (RNA-seq)

Reads in the Fastq format were quality controlled using fastqc and subsequently aligned to the *Mus musculus* GRCm39 reference genome (Ensembl). Reference genome was indexed using the Subread package (version 2.0.6) with paired-end settings. Adapters were trimmed using trimmomatic prior to alignement. Aligned reads were counted using featureCounts part of the Subread package (version 2.0.6). Count files were imported into R and quality controlled using RNASeqQC in R (version 0.1.4). Genes with a count sum below 20 across all samples were removed prior to analysis. We performed PCA on the samples to assess for outliers and underlying batch effects. Differential gene expression was done using DESeq2 in R (version 1.38.3) using default parameters, adding in batch coefficients to the model where required. An adjP < 0.05 threshold was set for calling significance, and volcano plots were generated using ggplot2.

#### Quartiling RNA and ChIP signal

To understand the relationship between H2A.Z and gene expression, raw RNA counts from each group were processed and quartiled using a custom R script. Briefly, RNA counts were normalized using the vst function in R (DESeq2), a row-mean normalized count value was determined and the gene set was split into even quartiles. H2A.Z signal was derived from the replicate merged BigWig files in deepTools. The H2A.Z signal was plotted on the quartiled mean gene expression data of each condition, respectively, using a custom R script. Statistics between 5xFAD H2A.Z signal and WT H2A.Z signal was computed using a Wilcox test in R (base R) by taking the signal distribution proximal to the TSS or distal to the TSS, and upstream or downstream the TSS, respectively.

#### ChIP signal on DEGs

To compute the change in H2A.Z signal on the genes changing with H2A.Z depletion, genes that either decreased, or increased in expression were subset. H2A.Z signal from the replicate merged BigWig files were then plotted at the TSS of these genes. Statistics were calculated using a Wilcox test in R.

### Stereotaxic surgery for viral delivery

H2A.Z.1 was depleted using a validated shRNA against *H2az1* packaged in AAV-DJ vector [14–16]. Mice were anesthetized with isoflurane and secured in a Kopf stereotaxic apparatus at 2.5 months of age. AAV viral particles were bilaterally injected into the CA1 region of the hippocampus at the following coordinates: anterior/posterior (AP) −2.0 mm, medial/lateral (ML) ±1.5 mm, and dorsal/ventral (DV) −1.6 mm. A total volume of 1 μL per hemisphere was delivered at a rate of 225 nL/min.

### Object location memory (OLM)

OLM was conducted during the early to mid-light phase, between 2 to 5 hours after lights on, in mice aged 3 to 8 months. The test arena consisted of a white melamine box (45 cm × 45 cm × 45 cm) placed in a room, with spatial cues presented on the walls in colors within the mice’s visible spectrum. Each mouse was brought from the holding room into the testing room and allowed to acclimate to the environment for 3 minutes before being placed in the test apparatus. The test consisted of three days. On Day 1 (habituation trial), each mouse was gently handled by the experimenter and placed in the empty test arena for 5 minutes. This trial was conducted only at 3 months of age. On Day 2 (training trial), mice were placed in the test arena with two identical novel objects (similar in size to the test mice), positioned in the two corners along the same wall, and allowed to explore for 10 minutes. On Day 3 (recall trial), one object remained in its original location, while the other was relocated diagonally to the opposite corner of the arena. Mice were then given 5 minutes to explore both objects. To minimize desensitization from repeated testing, a different set of objects was used each month, varying in color, shape, and material. The test arena and objects were cleaned with 70% ethanol between sessions to eliminate olfactory cues. All sessions were recorded with a digital camera mounted at the ceiling above the test arena, and later manually scored using EthoVision XT 17.5 software. Spatial memory performance was evaluated by comparing the time spent exploring the object in the novel location versus the familiar location and quantified as the discrimination ratio [DR = (seconds spent in novel location exploration – familiar location exploration)/(total seconds in object exploration)].

### Insoluble/soluble fractionation for pathology

GFP-expressing hippoamcpal tissue was dissected under a fluorescent lamp. Tissue was dounced in 200 µL of lysis buffer (1% Triton X-100 in 1× PBS supplemented with protease inhibitors) and transferred into a 0.2 mL Open Top Thickwall Cellulose Propionate Tube (Product #342303) and placed in an ultracentrifuge. Samples were spun at 100,000×g for 1 hour at 4°C, with acceleration/deceleration set to 4. The supernatant was collected as the soluble fraction and further diluted with an additional 200 µL of lysis buffer. The pellet was resuspended in 200 µL of 5 M guanidine HCl in 50 mM HCl (pH 8.0) with protease inhibitors. This lysate was then cleared by centrifugation at 20,000 × g for 20 minutes, and the insoluble fraction was collected. Protein concentrations for each fraction were measured using the Pierce™ Dilution-Free™ Rapid Gold BCA Protein Assay (Catalog #A55860).

### ELISA for Aβ and pTau-217

To measure Aβ_42_ levels, we used the Human Amyloid Beta 42 ELISA Kit, Ultrasensitive (KHB3544) according to the manufacturer’s instructions. The soluble fraction was diluted 1:3, and the insoluble fraction was diluted 1:100,000 using the Standard Diluent Buffer supplemented with 1 mM AEBSF before being loaded into the ELISA reaction. Phosphorylated tau (p-tau 217) was quantified using the PathScan® RP Phospho-Tau (Thr217) Sandwich ELISA Kit (#59672) according to the manufacturer’s protocol. The soluble fraction was diluted 1:200, and the insoluble fraction was diluted 1:800 using 1× Cell Lysis Buffer supplemented with 1× protease inhibitors before being loaded into the ELISA reaction. Aβ_42_ and pTau217 levels were normalized to protein concentration and further normalized to the Scramble control group for comparison with the H2A.Z.1-depleted group.

## RESULTS

### H2A.Z expression increases in the hippocampus of female, but not male AD patients

To determine if H2A.Z is dysregulated in human AD patients, we compared expression of two H2A.Z-encoding genes, *H2AZ1* (encodes H2A.Z.1) and *H2AZ2* (encodes H2A.Z.2), in post-mortem hippocampus. Expression of both genes increased in female, but not male AD patients (Figure 1A), demonstrating a sex-specific H2A.Z upregulation in AD. To determine if sex differences in expression are reflected in altered H2A.Z chromatin occupancy, we assessed H2A.Z binding using ChIP-sequencing (ChIP-seq) in post-mortem hippocampus from male and female AD patients and age-matched controls.

**Figure 1.**
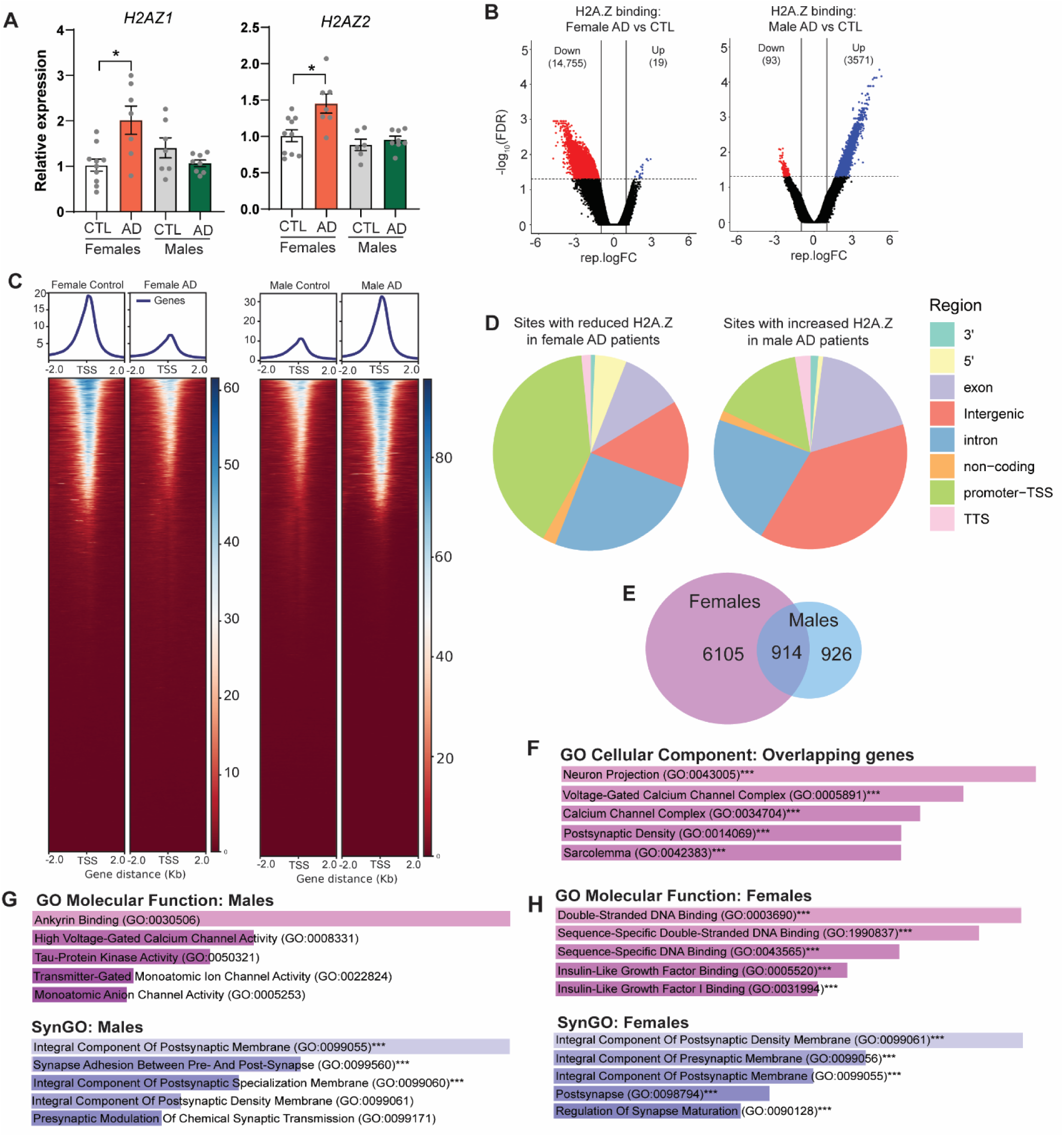
AD is associated with opposite shifts in H2A.Z binding in male and female patients. (A) qPCR analysis from hippocampus shows increased expression of H2A.Z encoding genes *H2AZ1* (Sex X AD Interaction: F_1,28_=12.19, p=0.002) and *H2AZ2* (Sex X AD Interaction: F_1,27_=4.26, p=0.049) in female compared to male AD patients (all post-hoc p<0.05). (B) H2A.Z ChlP sequencing ‘reveals opposing changes in female and male AD patients, whereby H2A.Z binding primarily declines in female (left) and increases in male (right) AD patients. (C) Heat map showing H2A.Z binding around the TSS. (D) Pie charts showing genomic distribution of differentially bound regions in females (left) and males (right). (E) Venn diagram showing overlap in DBGs in male and female patients. (F) Enrichr ontology for overlapping DBGs. (G) Enrichr gene ontology for differentially bound genes (DBGs) in female patients vs controls. (H) Enrichr gene ontology for DBGs in male patients vs controls.

### Male and female AD patients exhibit opposite changes in H2A.Z chromatin occupancy

H2A.Z binding showed opposite patterns of dysregulation in male and female AD patients, with more extensive changes occurring in women than men. Specifically, H2A.Z binding predominantly **decreased** in female (decreased at 14,755 loci and increased at only 19 loci) and **increased** in male AD patients (decreased at 93 loci and increased at 3,571 loci) (Figure 1B-C). These changes occurred on distinct genomic regions, whereby differentially bound regions (DBR) in females were predominantly localized to promoters, introns, exons, and 5’UTR, and male DBRs were primarily localized to intergenic sites, followed by introns, exons and promoters (Figure 1D). Notably, only female patients had altered H2A.Z binding on genes encoding *APP* (amyloid precursor protein) and *MAPT* (microtubule-associated protein tau), suggesting that H2A.Z may regulate pathology particularly in females.

H2A.Z changes occurred largely on distinct genes in male and female patients, with only 914 genes overlapping between sexes (Figure 1E). These common genes were enriched for terms related to neuronal signaling, including neuron projection, post-synaptic density, and calcium channels, indicating that H2A.Z may regulate synaptic function and neural activity in both sexes (Figure 1F). Indeed, differentially bound genes (DBGs) in male patients were enriched only for synaptic terms (Figure 1G), whereas female DBGs were also enriched for DNA-binding terms that encompassed diverse transcription factors, such as homeobox genes, SOX, GATA and FOX genes; zinc finger proteins, and bHLH factors, including *NEUROD2*, which is strongly implicated in cognitive function [27]. In females, H2A.Z was also enriched on several hormone receptors, most notably *NR3C1*, which encodes the glucocorticoid receptor, and *ESR1*, which encodes the estrogen receptor, suggesting that altered H2A.Z binding may regulate transcriptional programs throughout the female AD brain (Figure 1H).

### H2A.Z binding undergoes sex- and age-specific changes in 5xFAD mouse model of AD

We next utilized the well-established 5xFAD mouse model of AD to investigate the functional relevance of altered H2A.Z binding in human patient brains. We first conducted H2A.Z ChIP-seq to determine if sex-specific H2A.Z dysregulation observed in human AD tissue is recapitulated in 4-month-old 5xFAD mice. As with human patients, there was a dramatic sex difference, whereby extensive changes in H2A.Z binding occurred only in female mice. However, in contrast to human patients, 5xFAD females had a large **increase** rather than a **decrease** in H2A.Z binding, while males did not differ from controls (Figure 2A,B). Given that female AD patients lost and male AD patients gained H2A.Z, we next measured H2A.Z binding in 8-month-old mice to determine if mice at more advanced disease stage better recapitulate H2A.Z dysregulation in human disease. Indeed, H2A.Z binding declined dramatically in 5xFAD females at 8 months of age, reflecting a complete reversal of differential H2A.Z binding compared to the early disease stage (Figure 2D). As in human patients, male 5xFAD mice had increased H2A.Z binding at 8 months of age and these changes were much less extensive than in females (Figure 2C). Age related shifts in H2A.Z binding were further evidenced by comparing log fold change (logFC) on H2A.Z-bound loci at 4 and 8 months of age, whereby a clear shift from early accumulation to late depletion was evident only in female 5xFAD mice (Figure 2E,F).

**Figure 2.**
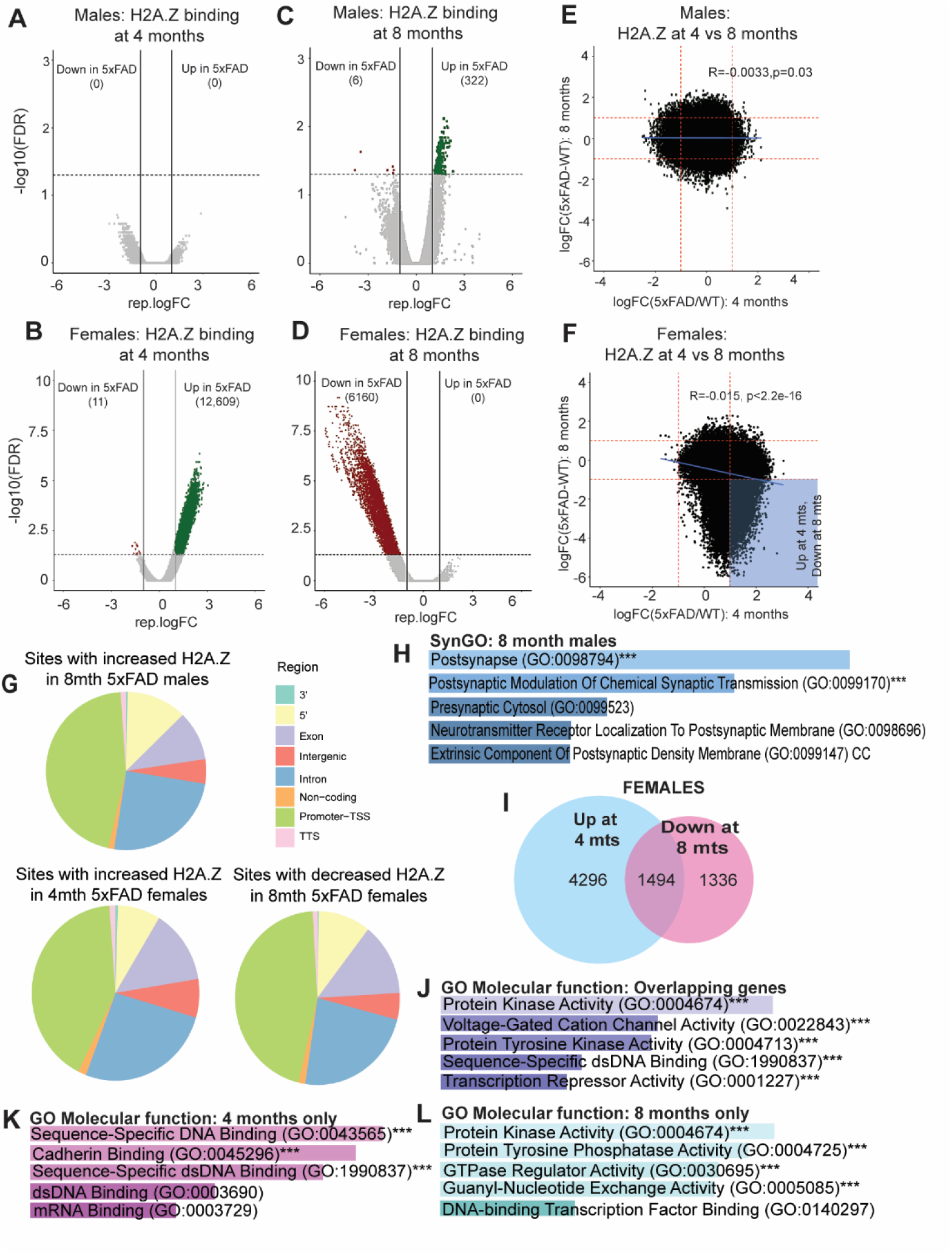
5xFAD mice exhibit sex- and age-specific shifts in H2A.Z binding. (A) In male mice, H2A.Z binding in 5xFAD mice did not differ from WT controls at 4 mts of age, but (C) increased at 8 mts of age. (B) In females, H2A.Z binding increased at 4 mts. and (D) decreased at 8 mts. in 5xFAD relative to WT controls. (E) Overall, there was no relationship between differentially bound genes at 4 and 8 mts. in males, whereas (F) females exhibited sites that increased binding at 4 mts. and decreased binding at 8 mts (lower right quadrant, shaded blue). (G) Genomic distribution of differentially bound sites is shown in pie charts. (H) Gene ontology (Enrichr) analysis of differentially bound genes (DBG) in 8 mth. old male mice. (I) Venn diagram showing overlap between DBGs modified at at 4 and 8 mts. (J) Gene ontology (Enrichr) for overlapping DBGs at 4 and 8 mts. (K), DBGs unique to 4mts and (L) DBGs unique to 8 mts.

In both sexes, differential H2A.Z binding localized primarily to promoters, introns and exons (Figure 2G), as in human female patients. Despite relatively few sites of H2A.Z accumulation in 5xFAD males, enrichment of affected sites for post-synaptic terms was consistent with synaptic enrichment in human male patients (Figure 2H; Figure 1F,G). Thus, 5xFAD mice exhibit sex- and age-specific changes in H2A.Z binding that support the utility of this model for studying the functional relevance of H2A.Z dysregulation in human patients.

### H2A.Z binding undergoes age-related shifts on a subset of genes in female mice

Given the dramatic shift in H2A.Z binding over disease progression in female mice, we examined if these changes occurred on the same or distinct genes at 4 and 8 months of age. Most genes that lost H2A.Z at 8 months first gained H2A.Z at 4 months (53%; 1494 of 2830, Figure 2I) and are enriched for protein kinase activity, including several genes involved in tau phosphorylation (e.g., *Dyrk2*, *Akt3*, *Mapk8*, *Mapk9*, and *Taok1*) and transcriptional regulation, as in female AD patients (Figure 2J). Notably, transcription-regulating genes were also enriched in genes modified only at 4 months of age, suggesting that transcriptional regulators undergo early shifts in H2A.Z binding (Figure 2K). In contrast, terms unique to 8-month-old females were enriched for protein kinase and phosphatase activity, including several genes involved in tau phosphorylation and dephosphorylation (Figure 2L). Notably, many genes that were modified in female patients were also modified in AD model mice (3127 of 7012; Supplemental Figure 1), including several tau-related genes like *GSK3B*, a key kinase in tau phosphorylation, *MAPT*, which encodes Tau, and *PTPRS*, which may protect against tau phosphorylation [28, 29]. Thus, differential H2A.Z binding is associated with key genes involved in amyloid and tau pathology and disease progression in female patients and AD model mice. Together, these data suggest that H2A.Z is highly dynamic over disease progression and that reduced H2A.Z binding in female AD patients may be preceded by H2A.Z accumulation in early disease stages.

### H2A.Z.1 depletion improves memory and pathology in female and impairs memory and pathology in male 5xFAD mice

Having established 5xFAD mice as a strong model of human changes in H2A.Z binding, we investigated the functional relevance of H2A.Z in memory by depleting the H2A.Z encoding gene *H2az1* in area CA1 of the hippocampus at 2.5 months of age in WT and 5xFAD mice. Mice were tested for object location memory (OLM) starting at 3 months and ending at 8 months of age to capture memory across disease progression and shifts in H2A.Z binding (Figure 3A). First, we confirmed that *H2az1* depletion was effective throughout testing (Supplemental Figure 2). Deficits in OLM emerged at 3 months of age in female 5xFAD mice, whereas male 5xFAD mice had intact memory at this age (Figure 3B). H2A.Z.1 depletion reversed memory impairment in the OLM task in 5xFAD females as early as 3 months of age and continued to improve memory until 8 months of age (Supplemental Figure 3). Interestingly, H2A.Z.1 depletion actually impaired memory in 5xFAD males at 3 months of age (Figure 3B), suggesting that H2A.Z.1 may be protective against memory decline at early disease stages in male 5xFAD mice. However, once 5xFAD males began to exhibit memory deficits at 4 months of age, H2A.Z.1 depletion no longer impacted memory formation, except for increasing memory irrespective of genotype at 5 months of age (Supplemental Figure 6). Critically, H2A.Z.1 depletion did not impact memory in WT mice of either sex, suggesting that H2A.Z.1 depletion is specifically acting on disease-related adaptations.

**Figure 3.**
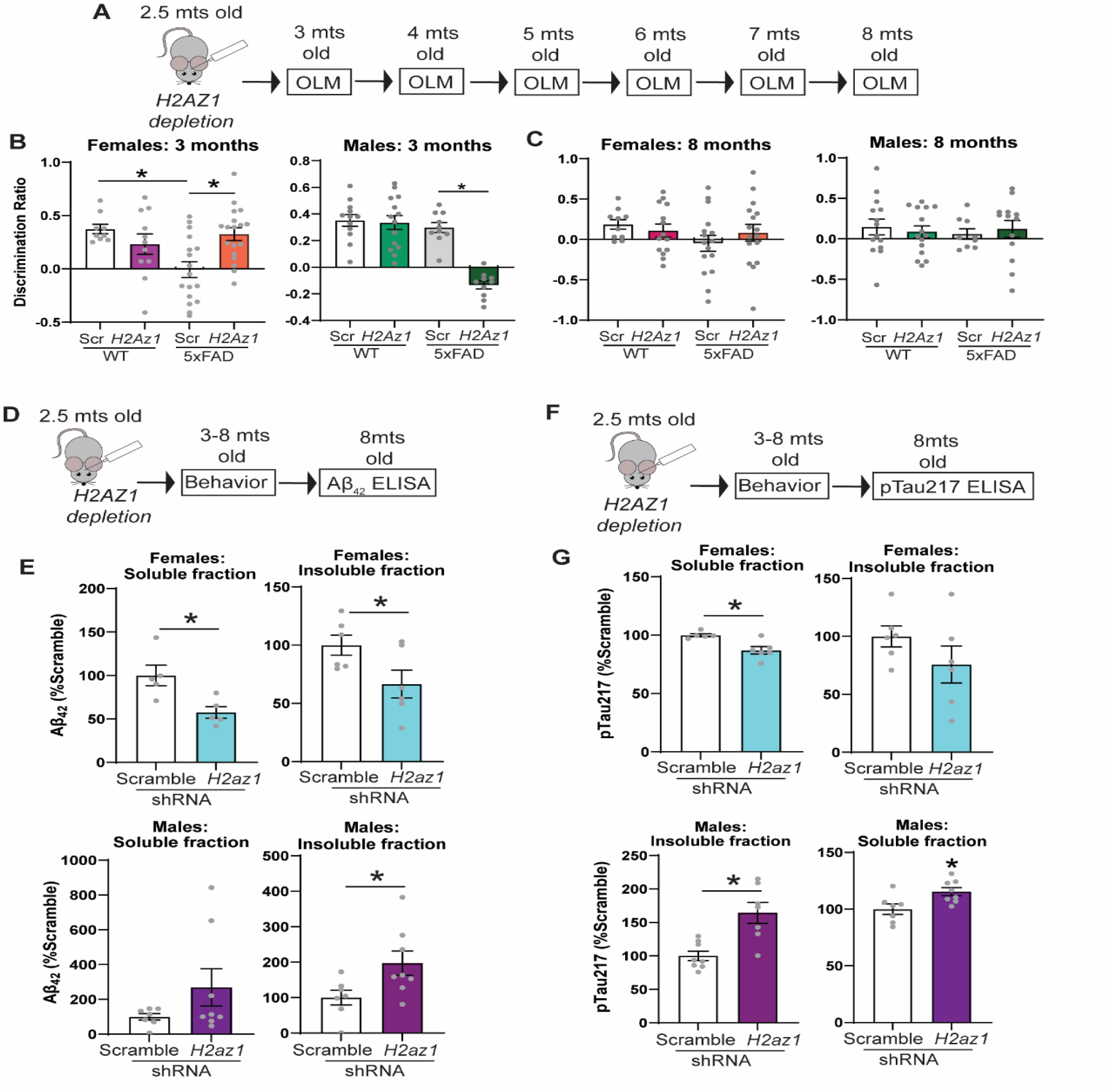
H2A.Z.1 depletion has opposite effects on memory decline and pathology in male and female 5xFAD mice. (A) Research design for behavioral testing. (B) At 3 months of age, *H2az1* depletion restored memory deficits in 5xFAD females (Genotype X Virus F_1,50_=9.09, p=0.004, *post-hoc p<0.05) and impaired normal memory in 5xFAD males (Genotype X Virus =20.91, p<0.001, *post-hoc p<0.05). (C) *H2az1* depletion had no impact on memory by 8 months of age. All intervening ages are showin in Supplemental Figure 6. (D) Research design for E. (E) *H2az1* depletion reduced soluble (ts=3.10, p=0.015) and insoluble (t10=2.27, p=0.047) 5xFAD is associated with distinct gene expression profiles in male and female mice

To determine if H2A.Z.1 depletion also regulates AD pathology, we measured soluble and insoluble Aβ_42_ and phosphorylated tau at threonine 217 (pTau217), which is a strong marker of AD pathology [30]. Consistent with behavioral data, H2A.Z.1 depletion reduced both soluble and insoluble Aβ_42_ in females and increased insoluble Aβ_42_ in males (Figure 3D-G). Similarly, H2A.Z.1 depletion reduced pTau217 in the soluble fraction in female mice and increased pTau217 in both fractions in male mice. Thus, H2A.Z.1 depletion is beneficial for memory and pathology in females and detrimental in males.

To elucidate the mechanism by which H2A.Z.1 depletion produces opposing effects on memory and pathology in male and female mice, we conducted RNA sequencing in infected hippocampal tissue of 4-month-old 5xFAD and WT mice. To determine if there are any pre-existing sex differences in disease-related gene regulation, we first compared effects of genotype (5xFAD *vs* WT) in mice injected with scramble control shRNA. In males, 5xFAD was associated with similar levels of up- and down-regulated gene expression, whereas gene expression was predominantly upregulated (81%) in 5xFAD female mice (Figure 4A). Only 43 DEGs (encoding lysosomal genes) were affected by 5xFAD in both males and females, indicating that disease-related gene dysregulation is sex specific (Supplemental Figure 4A,B).

**Figure 4.**
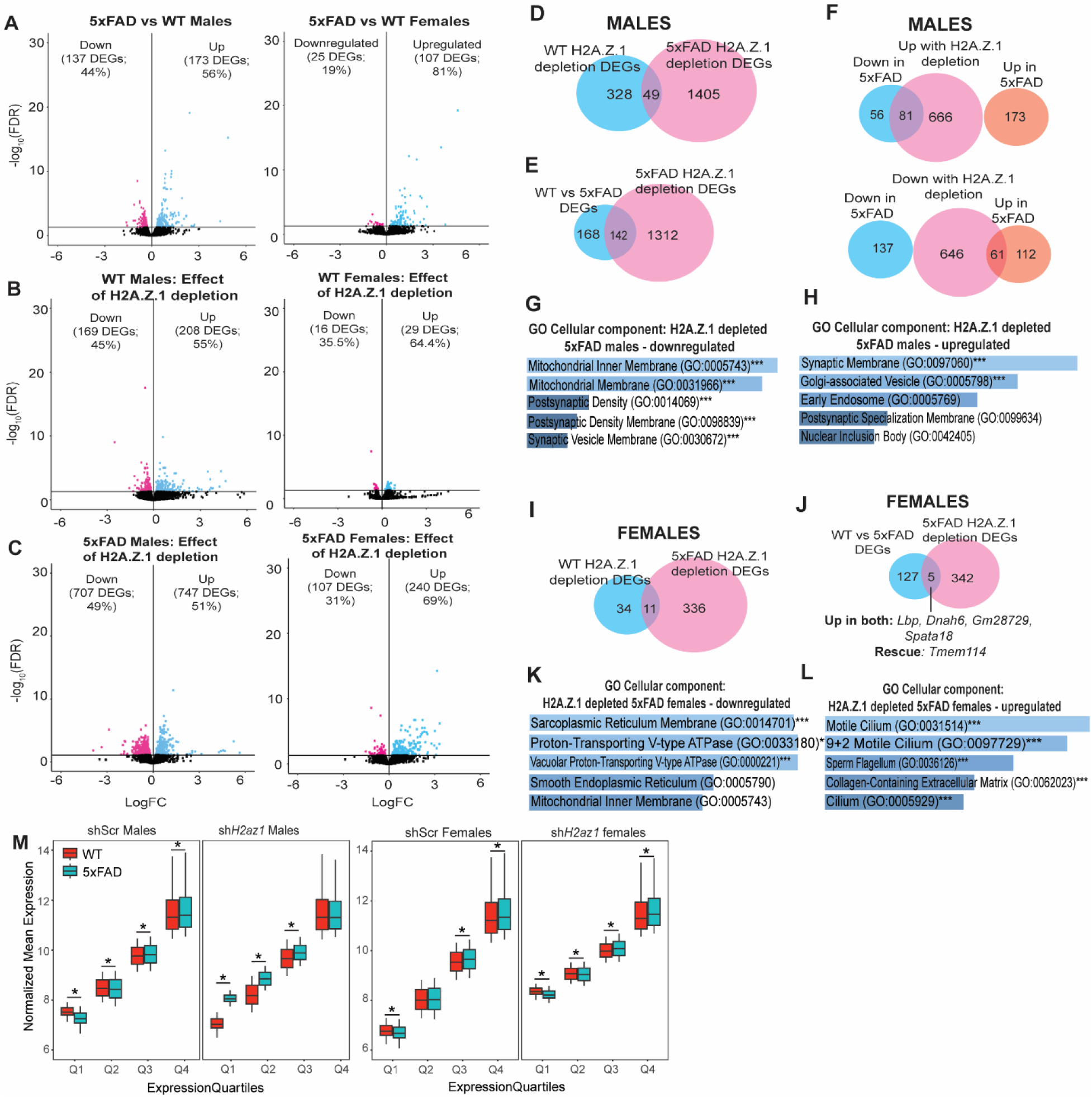
H2A.Z.1 depletion produces sex- and disease-specific effects on hippocampal gene expression. (A) Effect of genotype on gene expression in 5xFAD *vs.* WT scramble control males (left) and females (right). (B) Effect of *H2az1* depletion (vs. scramble control) in WT males (left) and females (right). (C) Effect of *H2az1* depletion (vs. scramble control) in 5xFAD males (left) and females (right). (D) Comparison of differentially expressed genes (DEGs) in response to *H2az1* depletion in WT and 5xFAD males (top). (E) Comparison of 5xFAD-related DEGs (5xFAD *vs.* WT) to DEGs altered by *H2az1* depletion in 5xFAD males (bottom). (F) Comparison of DEGs upregulated by *H2az1* depletion in 5xFAD males and DEGs that were either up- or down-regulated in 5xFAD *vs.* WT scramble control males (top). Comparison of DEGs downregulated by *H2az1* depletion in 5xFAD males and DEGs that were either up- or down-regulated in 5xFAD *vs.* WT scramble control males (bottom). (G) Gene ontology (Enrichr) for DEGs downregulated by *H2az1* depletion (vs. scramble control) in 5xFAD males. (H) Gene ontology (Enrichr) for DEGs upregulated by *H2az1* depletion *(vs.* scramble control) in 5xFAD males. (I) Comparison of DEGs in response to *H2az1* depletion in WT and 5xFAD females. (J) Comparison of 5xFAD-related DEGs (5xFAD *vs.* WT) to DEGs altered by *H2az1* depletion in 5xFAD females. (K) Gene ontology (Enrichr) for DEGs downregulated by *H2az1* depletion *(vs.* scramble control) in 5xFAD females. (L) Gene ontology (Enrichr) for DEGs upregulated by *H2az1* depletion *(vs.* scramble control) in 5xFAD females. **(M)** RNA expression in each quartile was compared between WT and 5xFAD males (left) and females (right) receiving scramble control AAV or H2A.Z.1 depletion. *adjP<0.05, Wilcox Test.

### H2A.Z.1 depletion has genotype- and sex-specific effects on gene expression

We next assessed effects of *H2az1* depletion and found a larger impact on gene expression in males than females irrespective of genotype, suggesting that H2A.Z.1 is a stronger transcriptional regulator in males (Figure 4C). Moreover, *H2az1* depletion had a bigger impact on gene expression in 5xFAD than WT mice irrespective of sex (Figure 4B-C), suggesting that H2A.Z.1 may be especially important for regulating transcription in disease. To elucidate sex-specific effects of H2A.Z.1 depletion in 5xFAD mice, we compared differentially expressed genes (DEGs) in each sex and found only 58 overlapping DEGs (Supplemental Figure 4C), consistent with divergent effects of *H2az1* depletion on memory and pathology. In 5xFAD males, H2A.Z.1 depletion impacted genes that were enriched for synaptic and ubiquitin-related terms, whereas the only enriched term in females was calcium ion binding and neuronal cilia (Supplemental Figure 4D), again indicating that H2A.Z contributes to AD through distinct pathways in each sex.

In males, *H2az1* depletion impacted distinct genes in WT and 5xFAD mice (Figure 4D), indicative of disease-specific roles in transcription. Genes that were downregulated with H2A.Z depletion in 5xFAD, but not in WT males were enriched for terms related to the mitochondrial membrane, post-synaptic density, synaptic vesicles, and potassium channels (Figure 4G). H2A.Z.1 depletion also reduced expression of genes relevant for amyloid pathology, including *Apbb1* (Aβ precursor protein binding family B member 1), a key intracellular adaptor involved in trafficking the Amyloid Precursor Protein (APP) and negatively regulating Aβ production [31]. Moreover, *Pink1* (PTEN induced putative kinase 1), whose loss is associated with Aβ accumulation and oxidative stress [32], was also downregulated by *H2az1* depletion, suggesting that increased Aβ levels with H2A.Z.1 depletion may be related to downregulation of genes that protect against amyloid accumulation in male mice.

Genes that were upregulated by *H2az1* depletion in 5xFAD males were enriched for synaptic membrane terms, including multiple GABA and glutamate receptors, suggesting that H2A.Z may contribute to altered synaptic function and excitability in AD [33]. Enriched terms also included the Golgi apparatus, whose dysfunction is implicated in Aβ formation [34] (Figure 4H). Notably, there was extensive upregulation of ubiquitin-related genes, including the E3 ubiquitin ligase *Itch* that is linked with cell death in amyloid-treated neurons and its upregulation may thus contribute to negative outcomes in *H2az1* depleted 5xFAD males [35]. Together, these data indicate that H2A.Z.1 may promote adaptive gene expression programs in AD and that negative outcomes of H2A.Z.1 depletion are associated with loss of this protective function.

To probe this hypothesis further, we investigated if H2A.Z.1 depletion impacts expression of disease-related DEGs (i.e., DEGs found in 5xFAD vs WT mice) and identified a subgroup of disease-modified genes that were “rescued” by *H2az1* depletion (e.g., increased in 5xFAD, decreased with H2A.Z.1 depletion; Figure 4E,F). These “rescued” genes were enriched for glutamate receptor activity and include the metabotropic glutamate receptor 8 (*Grm8*), which is elevated in 5xFAD mice and reduced by *H2az1* depletion. *Grm8* can protect cells from glutamate excitotoxicity, so its upregulation in 5xFAD males may be adaptive and disrupted by H2A.Z.1 depletion [36]. Similarly, the protective somatostatin receptor 2 (*Sstr2*) [37] is also upregulated with disease and downregulated by *H2az1* depletion, supporting a protective role of H2A.Z.1 in disease.

In contrast to males, H2A.Z.1 depletion in female mice only restored expression of one disease-regulated gene (*Tmem114*) gene and further increased expression of 4 upregulated genes, including *Lbp*, which has anti-inflammatory and neuroprotective effects [38] and *Spata18*, which has antioxidant functions and aids in DNA damage repair [39] (Figure 4I). However, most genes that were affected by H2A.Z.1 depletion were not altered by disease. Genes that were downregulated by H2A.Z.1 depletion in 5xFAD females were enriched for terms related to the endoplasmic reticulum, a key regulator of neuronal calcium associated with neurodegeneration in AD [40]. DEGs in this downregulated category include calcium regulators *Jph4* (junctophilin 4) and *Ryr1* (ryanodine receptor). RyR upregulation is associated with excessive calcium release and increased Aβ production, whereas RyR antagonism reduces Aβ pathology and improves memory in AD model mice, suggesting that H2A.Z.1-mediated *Ryr1* downregulation may contribute to beneficial effects of H2A.Z.1 depletion in female mice [41]. *Jph4* is also a critical calcium regulator that is highly expressed in the hippocampus [42], indicating that H2A.Z.1 depletion in females may improve memory and pathology by altering calcium regulation in the endoplasmic reticulum.

Top upregulated genes in *H2az1* depleted 5xFAD females included several candidates that are protective against AD, including *Ttr* (transthretin), a homo-tetrameric protein of therapeutic interest because it binds Aβ to reduce its toxicity and promote its clearance [43, 44]. Similarly, *A2m* (alpha-2-macroglobulin) promotes Aβ metabolism and alleles that reduce its expression increase risk of late onset AD [45], suggesting that elevated levels of these genes may reduce Aβ levels in 5xFAD females. In addition, gene ontology analyses show that upregulated genes are enriched for cilia-related terms (Figure 4J). Neuronal cells contain a single cilium that is critical for synaptic function and dysfunctional cilia are implicated in AD [46]. Thus, H2A.Z.1 regulates expression of multiple genes that may improve AD pathology and cognitive decline.

### H2AZ.1’s role as transcriptional repressor or activator depends on sex and disease

To better understand why H2A.Z.1 depletion has disease-specific effects on gene expression despite comparable levels of H2A.Z binding in WT and 5xFAD male mice, we sorted RNA-seq data into expression quartiles, with least expressed genes in quartile 1 (Q1) and most expressed genes in Q4. In male mice, H2A.Z.1 depletion enhanced disease-related differences in gene expression especially on genes binned in lower expression quartiles, where *H2az1* depletion reduced gene expression in WT mice and increased expression in 5xFAD mice (Figure 4M, Supplemental Figure 5A). These data suggest that H2A.Z.1 function shifts from primarily activating in WT to primarily repressive for lowly expressed genes in 5xFAD males, whereas impact on highly expressed genes was minimal. In contrast, *H2az1* depletion increased gene expression in both WT and 5xFAD females (Figure 4M, Supplemental Figure 5B), suggesting that H2A.Z.1 is more repressive to transcription in females than in males, irrespective of genotype. Together, these data suggest that despite minimal AD-related shifts in H2A.Z occupancy in male mice, loss of H2A.Z.1 extensively impacts gene expression in a disease-specific way.

### H2A.Z binding exhibits sex- and genotype-specific associations with gene expression

To assess the relationship between H2A.Z binding and gene expression, we compared H2A.Z ChIP signal on each expression quartile from scramble control mice. Consistent with our prior reports [14], there was a clear positive relationship between H2A.Z binding and gene expression in male WT mice, whereby H2A.Z binding increased with each increasing gene expression quartile (Figure 5A). This pattern was partly preserved in WT and 5xFAD females, who had increased H2A.Z binding from Q1 to Q2 but demonstrated overlapping H2A.Z signal at the top 2 quartiles. In 5xFAD males, H2A.Z binding exhibited lowest distinction between expression quartiles (Figure 5A), suggesting a disease-related shift in gene regulation.

**Figure 5.**
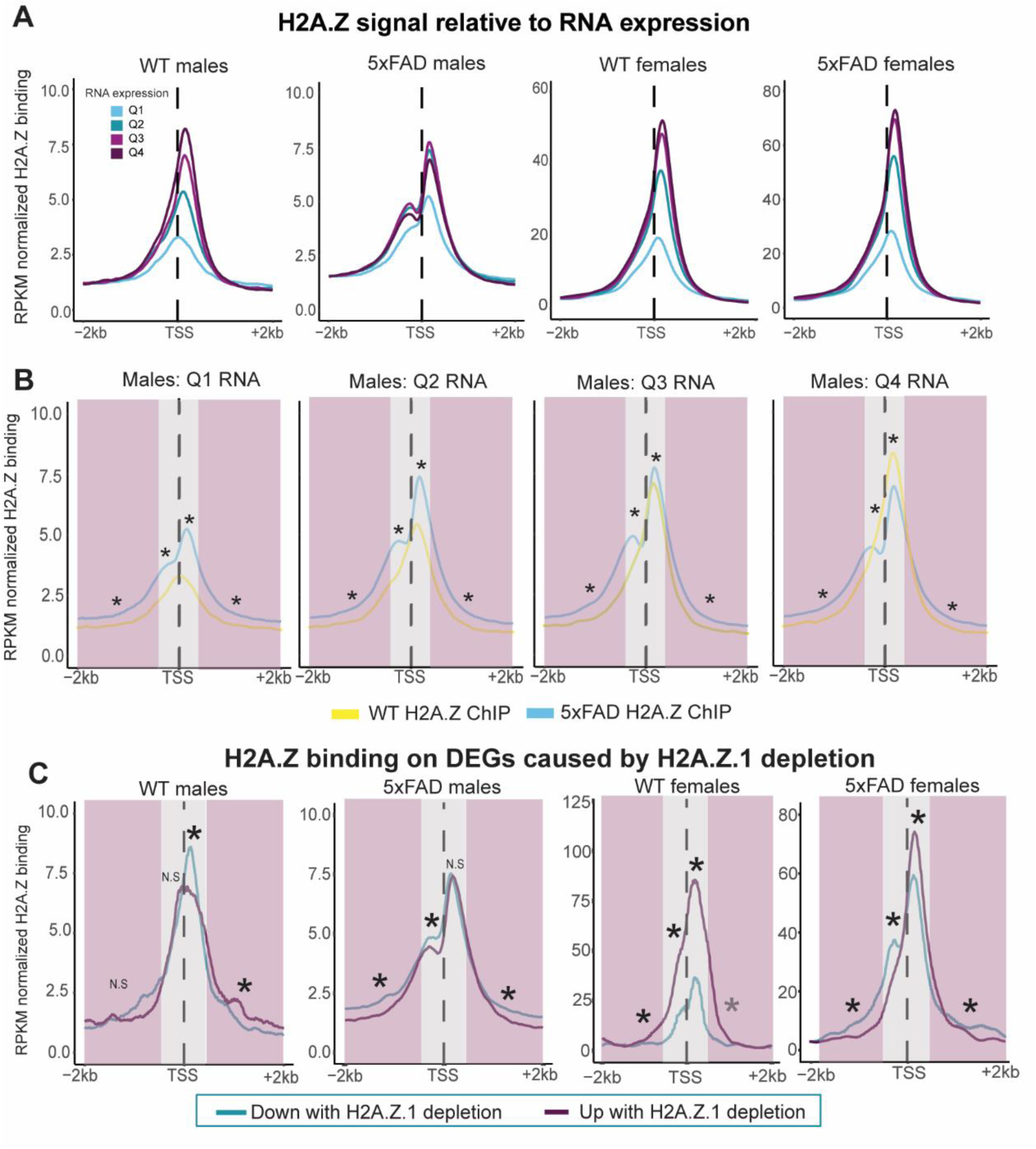
H2A.Z exhibits sex- and disease-specific associations with gene expression. (A) Gene expression was split into quartiles from least expressed (01) to most highly expressed (04) and H2A.Z levels (reads per kilobase per million; RPKM) were mapped for each quartile in WT and 5xFAD males (left) and females (right). (B) Data from A were replotted to show genotype effects in male mice at genes in different expression quartiles (01 =lowest expression, 04=highest expression. 5xFAD males had more expression at TSS-proximal and TSS-distal regions in 01-03. In 04, WT mice had more H2A.Z in TSS proximal regions, but less H2A.Z in TSS-distal regions *adj. p<0.05, Wilcox Test. (C) Normalized H2A.Z binding (RPKM) displayed for genes that were up- or downregulated with H2A.Z.1 depletion for WT and 5xFAD males (left) and females (right). adjP<0.05, Wilcox Test.

To better elucidate differential H2A.Z association with gene expression in males, we compared H2A.Z profiles between WT and 5xFAD mice at each quartile. 5xFAD males had more H2A.Z than WT mice at lower expression quartiles (Q1 and Q2) and less H2A.Z binding at the highest expression quartile (Figure 5B). Given that H2A.Z.1 depletion upregulated lowly expressed genes only in 5xFAD males (Figure 4M), these data suggest that H2A.Z accumulation takes on a repressive function in 5xFAD compared to WT males at less expressed genes.

To further clarify the relationship between H2A.Z binding and gene expression, we compared H2A.Z ChIP signal on genes that decreased vs. increased their expression with H2A.Z.1 depletion, assuming that depletion-induced repression indicates a positive transcriptional effect of H2A.Z in standard conditions. Prior work indicates that H2A.Z’s effect on gene activity is impacted by its upstream proximity to the TSS, whereby proximal promoter H2A.Z is more positively associated with gene expression than distal promoter H2A.Z [47]. In addition, intragenic H2A.Z downstream of the TSS is associated with reduced barrier to RNA polymerase compared to canonical H2A [48, 49], prompting us to compare H2A.Z binding on regions proximal (±350 bp from TSS) and distal (±350 to ±2000bp from TSS) to the TSS. We found distinct associations between H2A.Z and gene expression in WT and 5xFAD males. In WT males, proximal intragenic H2A.Z was more abundant on downregulated genes, consistent with a positive role in transcription. However, this relationship was not found for 5xFAD males, who instead had more H2A.Z on downregulated genes in all other positions (Figure 5C). Thus, even though total levels of H2A.Z binding do not change in male 5xFAD mice, the position of H2A.Z on genes sensitive to H2A.Z.1 depletion is redistributed, suggesting a disease-related shift in transcriptional regulation.

In contrast to males, WT females had more H2A.Z on up- than down-regulated genes at all positions, indicating a repressive role for H2A.Z on these genes (Figure 5C). This repressive function is consistent with gene de-repression in H2A.Z.1 depleted females (Supplemental Figure 5) and with more depletion induced up- than down-regulation only in females (Figure 4B,C), suggesting that H2A.Z has sex-specific transcriptional outcomes. As in WT females, 5xFAD females had more H2A.Z on up- than down-regulated genes immediately downstream of the TSS, but more H2A.Z on down- than up-regulated genes at all other positions, pointing to disease-related changes in H2A.Z distribution.

To elucidate why H2A.Z has distinct transcriptional functions in females, we examined its co-localization with AD-related modifications on other histones that can modify H2A.Z’s transcriptional outcomes [50]. Specifically, we compared H2A.Z levels on genomic regions that either lost or gained repressive (H3K27me3) or activating (H3K27ac; H3K4me3) histone marks in female CK-p25 mouse model of AD [26]. At 4 months, H2A.Z signal was enriched on sites that lost, but not sites that gained H3K4me3 (a marker of active promoters) in CK-p25 females. Moreover, H2A.Z was enriched on sites that gained, but not sites that lost the repressive H3K27me3 mark (Supplemental Figure 6), suggesting that H2A.Z is enriched particularly on regions that become repressed in female AD mouse models.

## DISCUSSION

Here, we provide first evidence for histone variant dysregulatoin in AD, thus opening a new avenue for understanding how disease-related shifts in chromatin contribute to transcriptional dysregulation, pathology, and memory decline. Specifically, we show that histone H2A.Z has sex- and disease-specific roles in gene regulation, memory, and pathology, whereby H2A.Z depletion is beneficial in females and detrimental in males. Importantly, these effects are disease specific, indicating that H2A.Z dysfunction is tapping into a core feature of AD rather than a general effect on memory formation.

The most striking observation is that H2A.Z binding changes in the opposite direction in male and female patients and that manipulating H2A.Z.1 results in opposite disease outcomes in male and female mice. Notably, different risk genes predict amyloid and tau pathology in post-mortem tissue of male and female AD patients, indicating that sex-specific pathways may contribute to pathology [4]. Our data point to several potential pathways by which H2A.Z may produce beneficial impacts in females and detrimental effects in males. For example, H2A.Z binding was modified on key pathology-related genes (*APP* and *MAPT*) in human female, but not male patients and mouse models, suggesting that shifts in H2A.Z binding at these genes may alter pathology outcomes. Moreover, H2A.Z.1 depletion produced expression changs that are consistent with improved outcomes only in female mice, including upregulation of *Ttr*, which promotes Aβ clearance [43, 44] and downregulation of ryanodine receptors that can promote Aβ production [41]. In contrast, H2A.Z.1 depletion in male mice reduced expression of several protective genes, including the *Apbb1* and *Pink1*, which may contribute to increased Aβ levels in males [31, 32]. Moreover, H2A.Z.1 depletion in males reversed several disease-related expression changes that may be adaptive in AD, including upregulation of *Grm8* and *Sstr2*, which are protective against excitotoxicity and neuronal damage [36, 37]. Thus, H2A.Z may be critical for maintaining an adaptive transcriptional profile in males and its depletion disrupts this adaptive function.

Another key observation was the temporal shift in H2A.Z binding over disease progression, whereby H2A.Z accumulation at early stages precedes H2A.Z depletion at late disease stages in female mice. A study of female CK-p25 mice also reported transient, persistent, and late-emerging epigenetic changes in various histone modifications with disease progression [26], indicating that the epigenome may be responsive to disease progression and in turn impacts disease progression through altered gene expression. Consistently, we observed that H2A.Z binding is altered on ion channels in both male and female AD patients (Figure 1F), which may relate to increased neuronal excitability early in disease and decreased excitability in later disease stages [51]. Notably, countering early H2A.Z accumulation with AAV vector mediated H2A.Z.1 depletion produced persistently beneficial effects on memory and pathology in female mice over 5 months, suggesting that interfering with early H2A.Z accumulation can modify the course of the disease.

Finally, our data suggest that H2A.Z’s role in transcription varies with sex and disease, whereby H2A.Z predominantly promotes gene activity in WT males but has a mix of repressive and activating effects in females and 5xFAD males. H2A.Z has a complex role in transcription that can be modified by various factors, including the number of H2A.Z histones present in a nucleosome, its positioning relative to the TSS, its post-translational modifications, and the factors it interacts with [13, 50]. Here, we find that genomic regions that accumulate repressive PTMs in mouse models of AD have more H2A.Z, suggesting that coupling of H2A.Z with repressive PTMs may account for more repressive transcriptional role in disease.

Overall, our data show that males and females exhibit divergent changes in H2A.Z binding in Alzheimer’s disease that are associated with distinct effects on transcription, memory, and pathology. The distinct behavioral and pathological outcomes with H2A.Z.1 depletion emphasize the need to investigate sex-specific contributions to AD to design sex-appropriate therapeutic interventions.

## Author contributions

JQL, LAH, SC, and IBZ designed the study, analyzed the data and co-wrote the manuscript. JQL, LAH, SC, TABM, TAD, ZJ, and FP conducted the experiments. BJW co-wrote the manuscript, helped with student supervision and AAV design. SMW, ILR, and MAB packaged AAV constructs. LAH, PJM and SH analyzed ChIP sequencing data. MF conducted RNA analysis in human tissue.

