## Supplemental Figure 1 for "Opposing effects of histone H2A.Z on memory, transcription and pathology in male and female Alzheimer’s disease mice and patients"

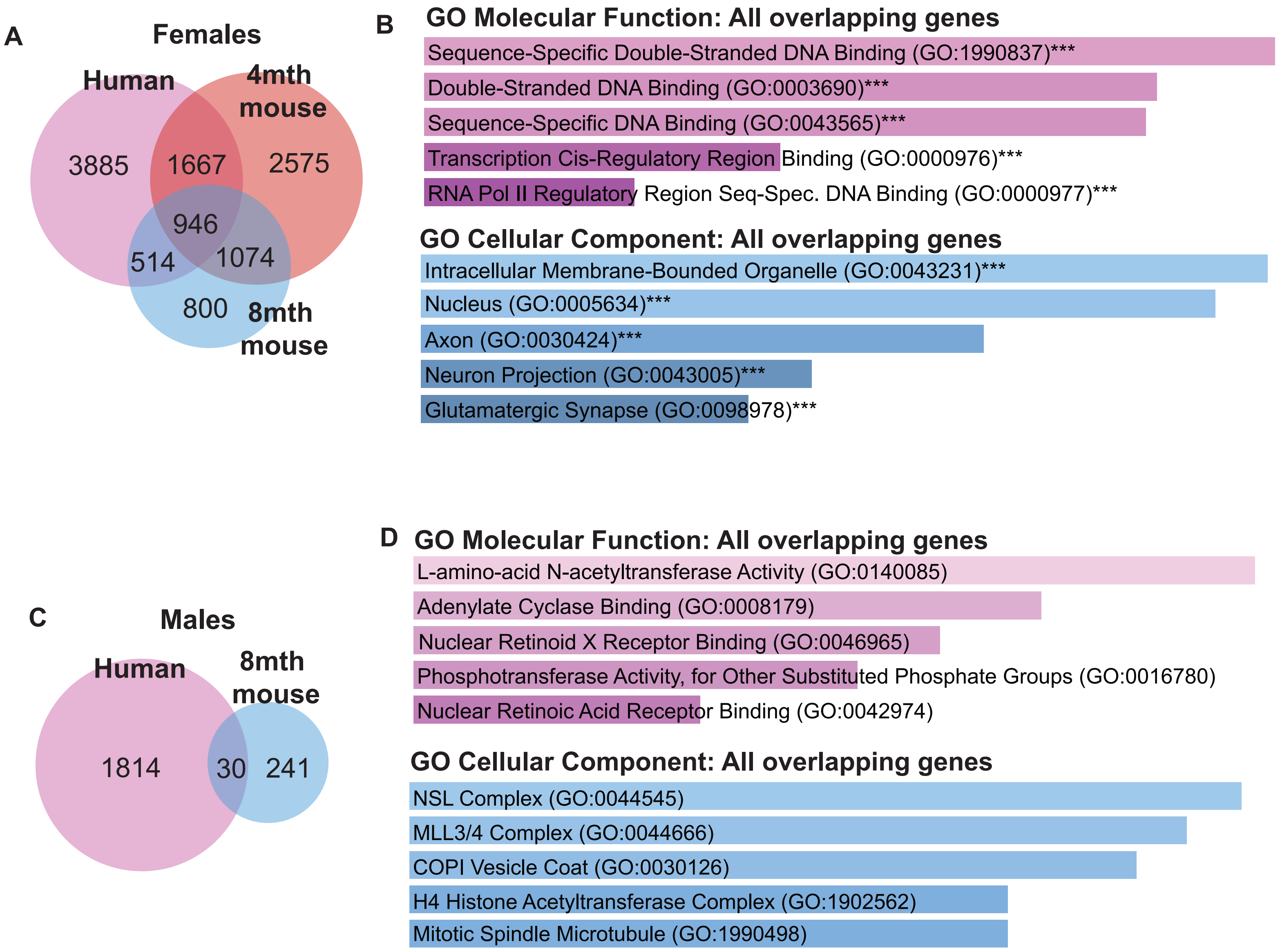


***Supplemental Figure 1. There is extensive overlap between genes with differential H2A.Z binding in human AD females and 5xFAD model mice.*** (A) Venn diagram depicting differentially bound genes (DBGs) in female AD patients and 5xFAD mice. (B) Gene ontology (Enrichr) for overlapping genes using GO Molecular Function (top) and GO cellular component (bottom). ***Indicates FDR<0.05. (C) Venn diagram depicting differentially bound genes (DBGs) in male AD patients and 5xFAD mice. (D) Gene ontology (Enrichr) for overlapping genes using GO Molecular Function (top) and GO cellular component (bottom).
