## Supplemental Figure 2 for "Opposing effects of histone H2A.Z on memory, transcription and pathology in male and female Alzheimer’s disease mice and patients"

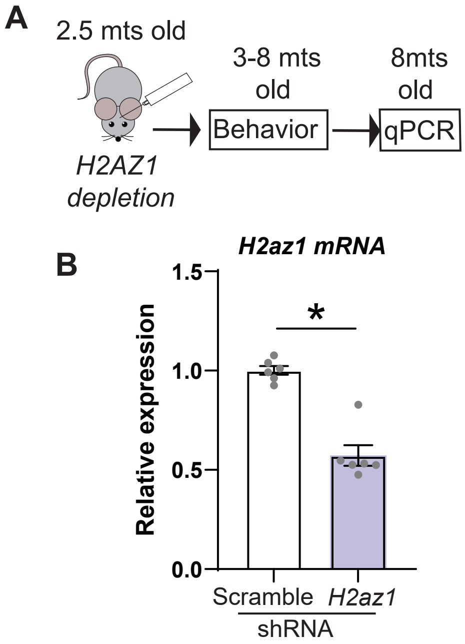


***Supplemental Figure 2. Validation of hippocampal H2az1 depletion***. (A) Research design for tissue collection. (B) *H2az1* shRNA reduced *H2az1* expression compared to scramble controls ($t_{10}$=6.96, $p<0.001$)
