## Supplemental Figure 3 for "Opposing effects of histone H2A.Z on memory, transcription and pathology in male and female Alzheimer’s disease mice and patients"

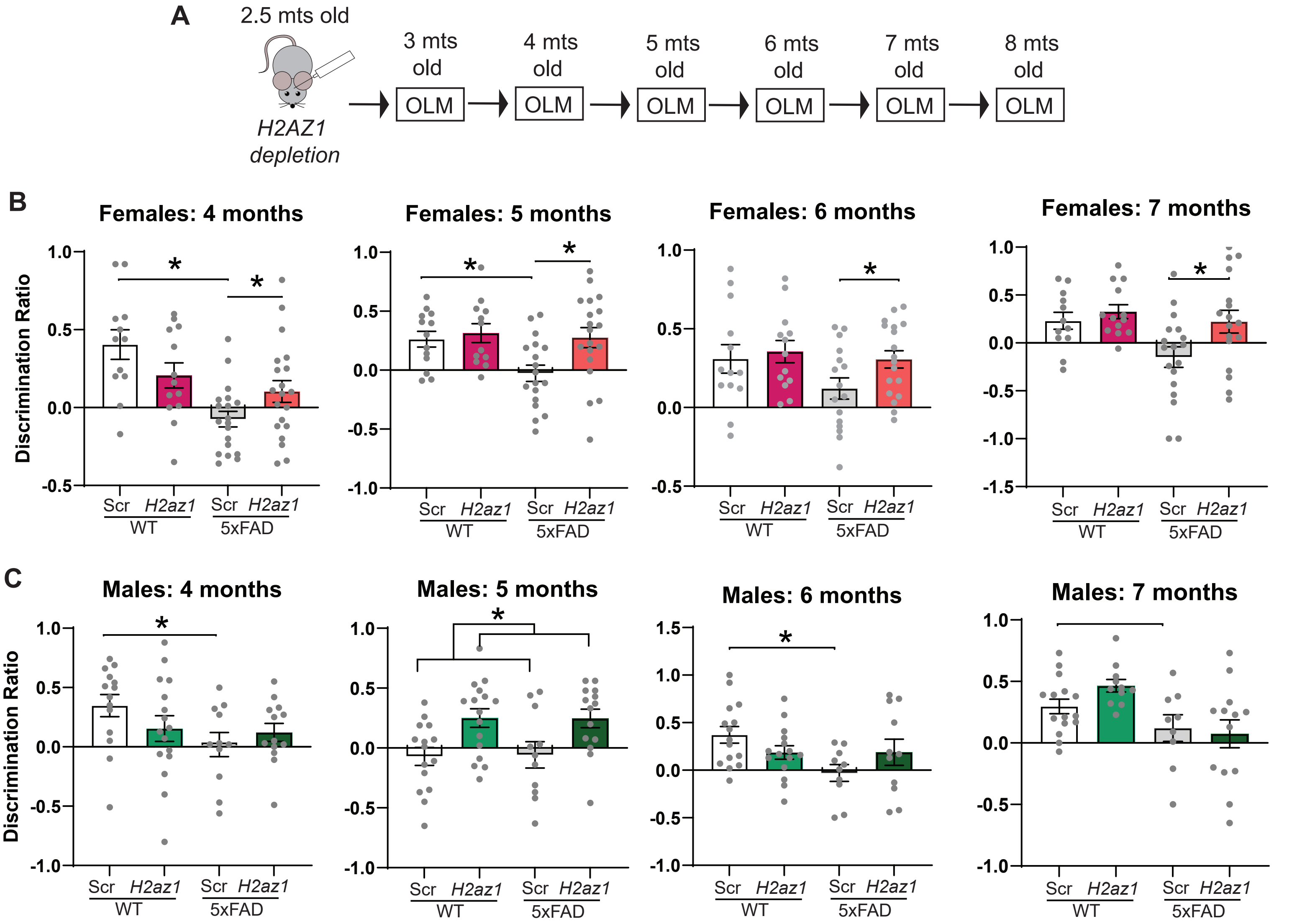


***Supplemental Figure 3. H2A.Z.1 has sex-specific effects on object location memory.*** (A) Study design for behavioral testing. Data for months 3 and 8 are presented in Figure 3. (B) 5xFAD females had improved memory with *H2az1* depletion at 4 (Genotype X Virus $F_{1,58}$=7.90, p=0.007, *post-hoc p<0.05), 5 (Genotype $F_{1,57}$=4.03, p=0.049, Virus $F_{1,57}$=4.57, p=0.04, *post-hoc p<0.05), 6 (*p<0.05 post-hoc only) and 7 (Genotype $F_{1,55}$=3.97, p=0.05, Virus $F_{1,55}$=4.44, p=0.04, *post-hoc p<0.05) months of age. (C) *H2az1* depletion did not impact OLM memory in 5xFAD males at any age, but there was a main effect of Virus at 5 months of age (Virus $F_{1,52}$=13.67, p<0.001), whereby memory improved in *H2az1* depleted mice irrespective of genotype. Deficits in 5xFAD vs. WT as indicated, *p<0.05.
