## Supplemental Figure 4 for "Opposing effects of histone H2A.Z on memory, transcription and pathology in male and female Alzheimer’s disease mice and patients"

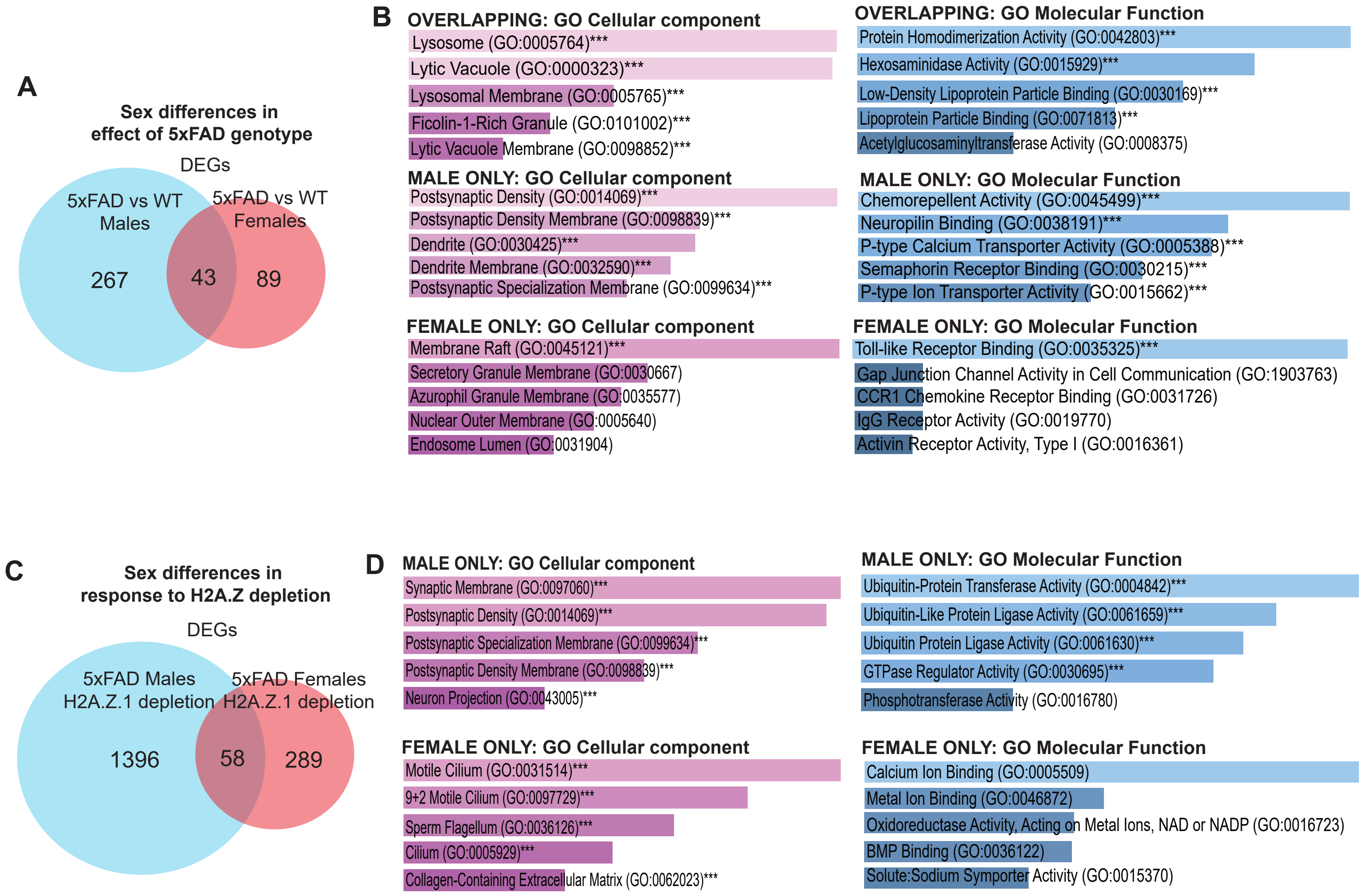


***Supplemental Figure 4. Male and female mice exhibit distinct profiles of gene dysregulation with 5xFAD genotype and H2A.Z.1 depletion.*** A) Venn diagram comparing differentially expressed genes (DEGs) produced by comparing 5xFAD and WT mice in the scramble control group for each sex. B) Gene ontology analyses (Enrichr) for DEGs that overlap between the sexes (top), for genes that are uniquely modified in males (middle) and genes that are uniquely modified in females (bottom). C) Venn diagram comparing DEGs produced by *H2az1* depletion in 5xFAD mice *vs.* scramble controls in each sex. D) Gene ontology analyses (Enrichr) for DEGs that are uniquely modified in males (top) and genes that are uniquely modified in females (bottom).
