## Supplemental Figure 5 for "Opposing effects of histone H2A.Z on memory, transcription and pathology in male and female Alzheimer’s disease mice and patients"

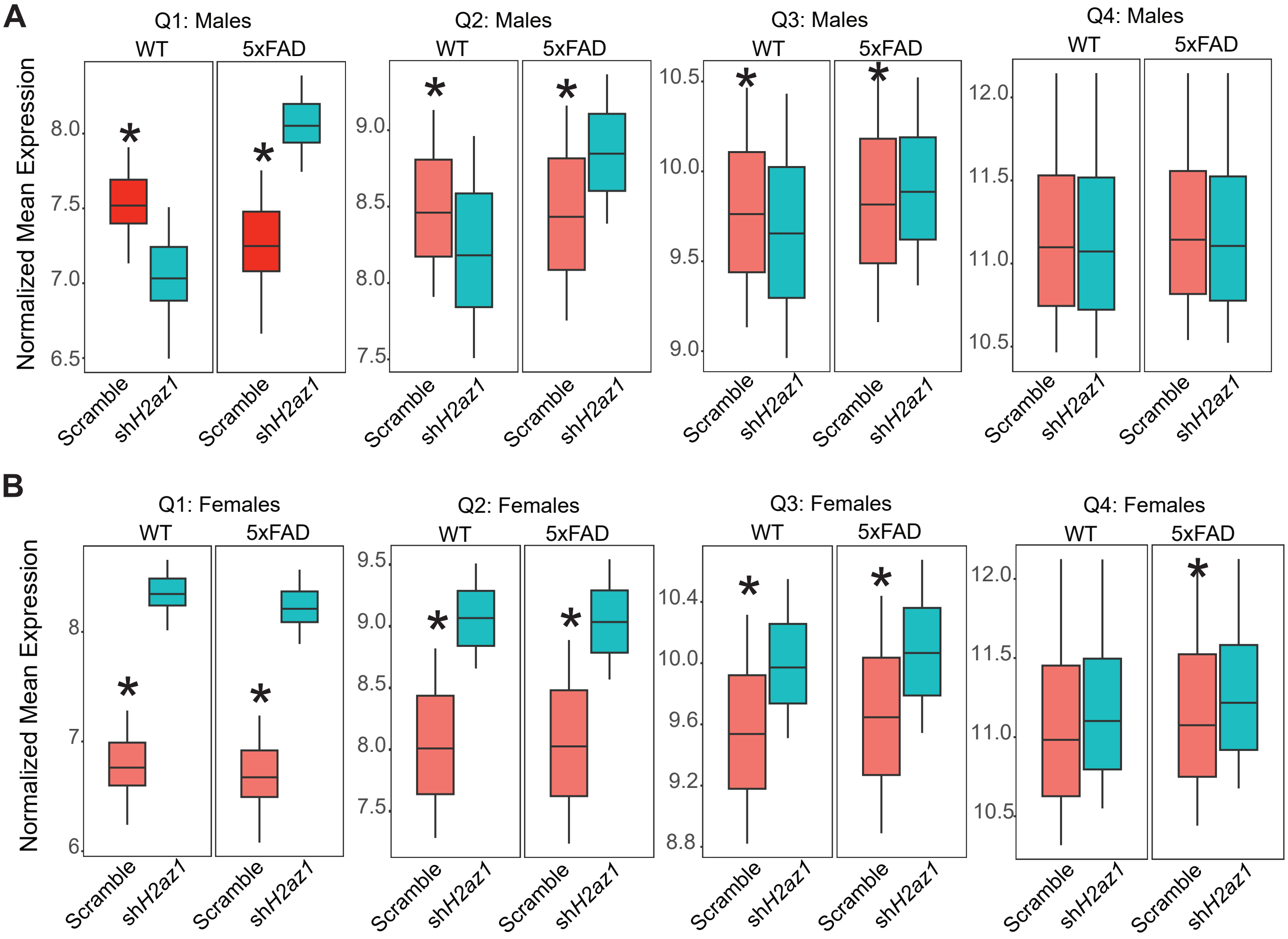


***Supplemental Figure 5. H2A.Z.1 depletion has genotype-specific effects on gene expression in male, not in female mice.*** Gene expression quartiles from Figure 4B were rearranged to more easily interpret effects of H2A.Z.1 depletion in WT and 5xFAD males (A) and females (B). *AdjP<0.05, Wilcox Test.
