## Supplemental Figure 6 for "Opposing effects of histone H2A.Z on memory, transcription and pathology in male and female Alzheimer’s disease mice and patients"

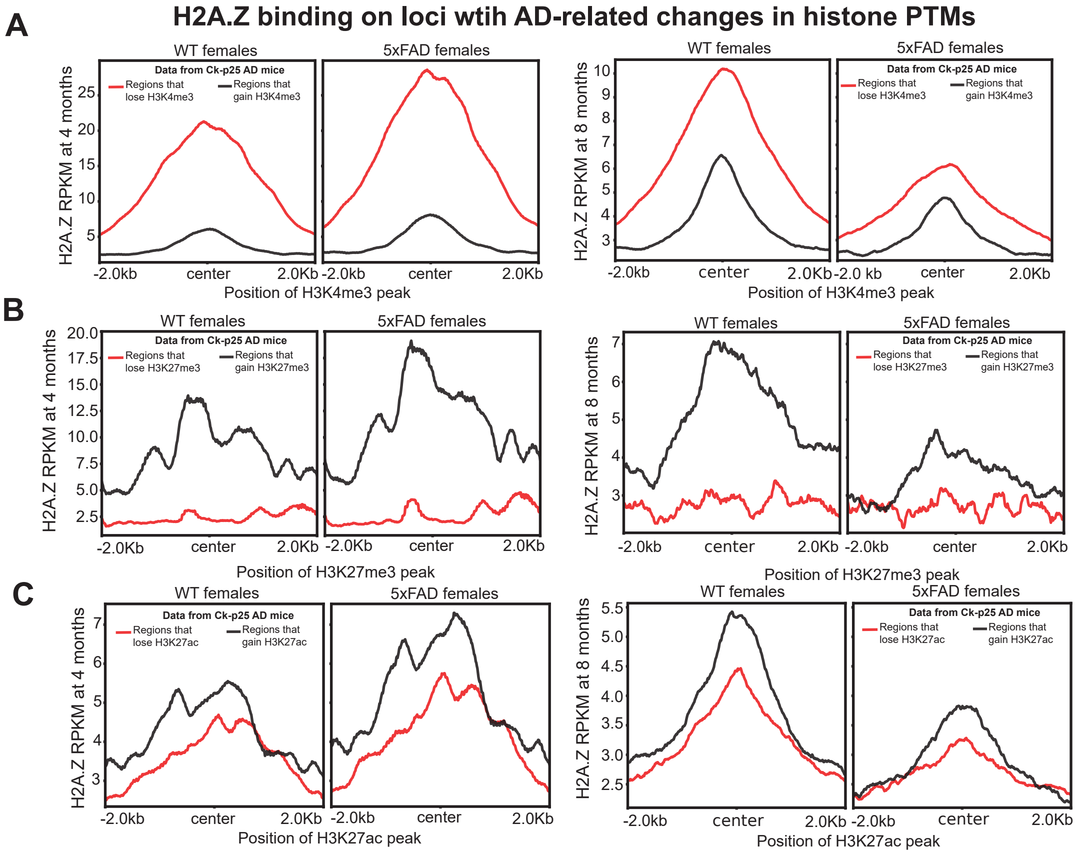


***Supplemental Figure 6. H2A.Z binding is enriched on regions that gain repressive, but not active histone H3 marks in CK-p25 mice.*** H2A.Z signal in 4 (left) and 8 (right) month old female mice was mapped on sites that gained or lost the activating mark H3K4me3 (A), the repressive mark H3K27me3 (B), or the activating mark H3K4me3 (C). Histone modifications were extracted from female Ck-p27 mice in Gjoneska et al (2015).
